# Switch from translation initiation to elongation needs Not4 and Not5 collaboration

**DOI:** 10.1101/850859

**Authors:** George E Allen, Olesya O Panasenko, Zoltan Villanyi, Marina Zagatti, Benjamin Weiss, Christine Polte, Zoya Ignatova, Martine A Collart

## Abstract

Not4 and Not5 are crucial components of the Ccr4-Not complex with pivotal functions in mRNA metabolism. Both associate with ribosomes but mechanistic insights on their function remain elusive. Here we determine that Not5 and Not4 synchronously impact translation initiation and Not5 alone alters translation elongation. Deletion of Not5 causes elongation defects in a codon-dependent fashion, increasing and decreasing the ribosome dwelling occupancy at minor and major codons, respectively. This larger difference in codons’ translation velocities alters translation globally and enables kinetically unfavorable processes such as nascent chain deubiquitination to take place. In turn, this leads to abortive translation and favors protein aggregation. These findings highlight the global impact of Not4 and Not5 in controlling the speed of mRNA translation and transition from initiation to elongation.

**Summary:** Not4 and Not5 regulate translation synchronously but distinguishably, facilitating smooth transition from initiation to elongation

## Results and discussion

The Ccr4-Not complex is a global regulator of mRNA metabolism in eukaryotic cells (for review see (1)). It regulates transcription and RNA quality control in the nucleus (2–4) and represses gene expression in the cytoplasm (5,6). Ccr4-Not plays an important role in co-translational processes. Not1, the largest subunit, assembles in granules, assemblysomes, protecting stalled ribosomes from the ribosome quality control (RQC) machinery and enabling co-translational association of interacting subunits of protein complexes (7). Not5 facilitates translation of transcripts encoding ribosomal proteins (RPs) through Not1-mediated mRNA binding (8), however mechanistic details are elusive. The Not4 RING E3 ligase ubiquitinates the Rps7 ribosomal protein (9) which, following ribosome stalling, signals co-translational mRNA degradation (NGD) when RQC is defective (10). The roles of Not4 and Not5 in translation and the mechanism of their action under permissive growth conditions is unknown.

To characterize the impact of Not4 and Not5 in translation we performed cell-wide analysis of translation using ribosome profiling (11) in Not4- (*not4Δ*) and Not5-depleted (*not5Δ*) strains and compared it to wild-type yeast (7) under permissive growth conditions. In total, 5048 transcripts were detected as genuinely translated above the threshold (reads per kilobase per million of sequencing reads (RPKM) >1, **Table S1**), with very good reproducibility between biological replicates (**Fig. S1A**), well-defined three-nucleotide periodicity (**Fig. S1B**) and the majority of ribosome protected fragment (RPF) lengths between 28 and 31 nucleotides (**Fig. S1C**). Metagene analysis of the aggregated RPFs across transcripts (12) in *not4Δ* showed a significant RPF accumulation at the start codon with no impact on elongation (**Fig. 1A**, upper panel). In *not5Δ*, along with the increased RPFs at start, we observed a marked RPF increase within the first 100 codons at the 5’ end of coding sequences (CDS), followed by a relaxation of the coverage relative to that of the wild type (**Fig. 1A**, lower panel). Together with a higher ratio between RPFs at the beginning and end of each CDS in *not5Δ* compared to wild type, particularly for high expressed genes (**Fig. 1B**), these observations suggest defective translation elongation in *not5Δ*.

**Fig. 1.**
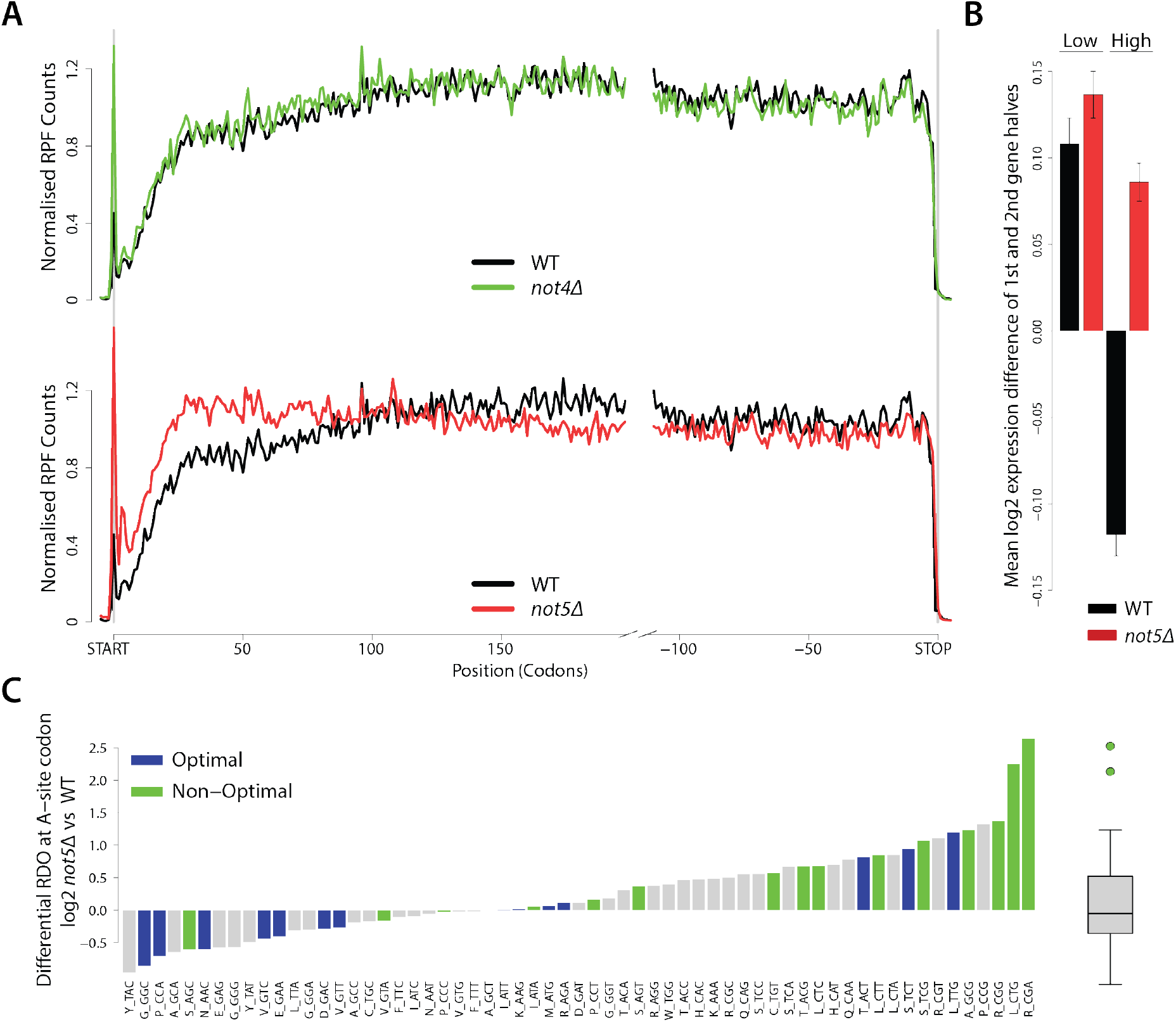
Translation elongation is altered in *not5Δ*. (**A**) Metagene analysis in wild-type, *not4Δ* (top) and *not5Δ* (bottom) strains. (**B**) Mean RPKM ratio of both halves of ORFs in *not5Δ* and wild type, split by high and low expression (greater or lesser than median expression). SEM error bars. (**C**) RDO at A-site codons in *not5Δ* relative to wild type, with 15 most (blue) and 15 least used (green) codons. Grey, all other codons. Optimal codons are enriched in the lower RDO group *not5Δ* (proportion test; p = 0.01127; logFC<−0.2) relative to higher (logFC>0.2) and non-optimal codons are enriched in the higher RDO group (p = 0.04327).

To understand the elongation defect in *not5Δ*, we determined the ribosome dwelling occupancy (RDO) at each codon in the ribosomal A and P sites (that are the binding sites of aminoacyl-tRNA and peptidyl-tRNA, respectively), by calibrating the 5’ ends of RPFs to the start codon (13). RDO correlates with the speed of translation of a codon, with slow translated codons exhibiting higher RDO than fast translated ones (14). The RDO at the A site codons in *not5Δ* was markedly altered, with a significant reduction at optimal and increase at non-optimal codons (**Fig. 1C**). The changes in the RDO in the P site were less prominent, without any correlation to codon usage (**Fig. S1D**). Notably, the changes in A site RDO in *not5Δ* were the same for transcripts with low and high expression (**Fig. S1E**), despite the skewed distribution of optimal and non-optimal codons between high and low expressed transcript groups (**Fig. S1F**).

The concentration of the cognate aminoacyl-tRNA is one major determinant of the ribosome speed at a codon in the ribosome A-site. We measured the concentrations of total and aminoacylated tRNAs using tRNA-tailored microarrays. The overall expression of tRNAs was similar in *not5Δ* and wild type with exception of few tRNAs up-regulated (e.g. two tRNA^Ser^, one tRNA^Gly^ and one tRNA^Asp^) and down-regulated (one tRNA^lys^, one tRNA^Gln^, one tRNA^Cys^ and one tRNA^Thr^) (**Fig. 2A**, top and **Table S2**). To our surprise, the overall amount of aminoacyl-tRNAs was even higher in *not5Δ* (**Fig. 2A**, bottom and **Table S2**) and correlated with a lower concentration of free amino acids in *not5Δ* (**Fig. 2B**). The altered A site RDO in *not5Δ* did not correlate with cognate tRNA paucity. The global accumulation of aminocyl-tRNAs that are not used in translation was rather consistent with the overall reduction of translation elongation (**Fig. 2C**). Hence, changes in RDO at A site codons was clearly dependent on Not5 and not on tRNA amounts.

**Fig. 2.**
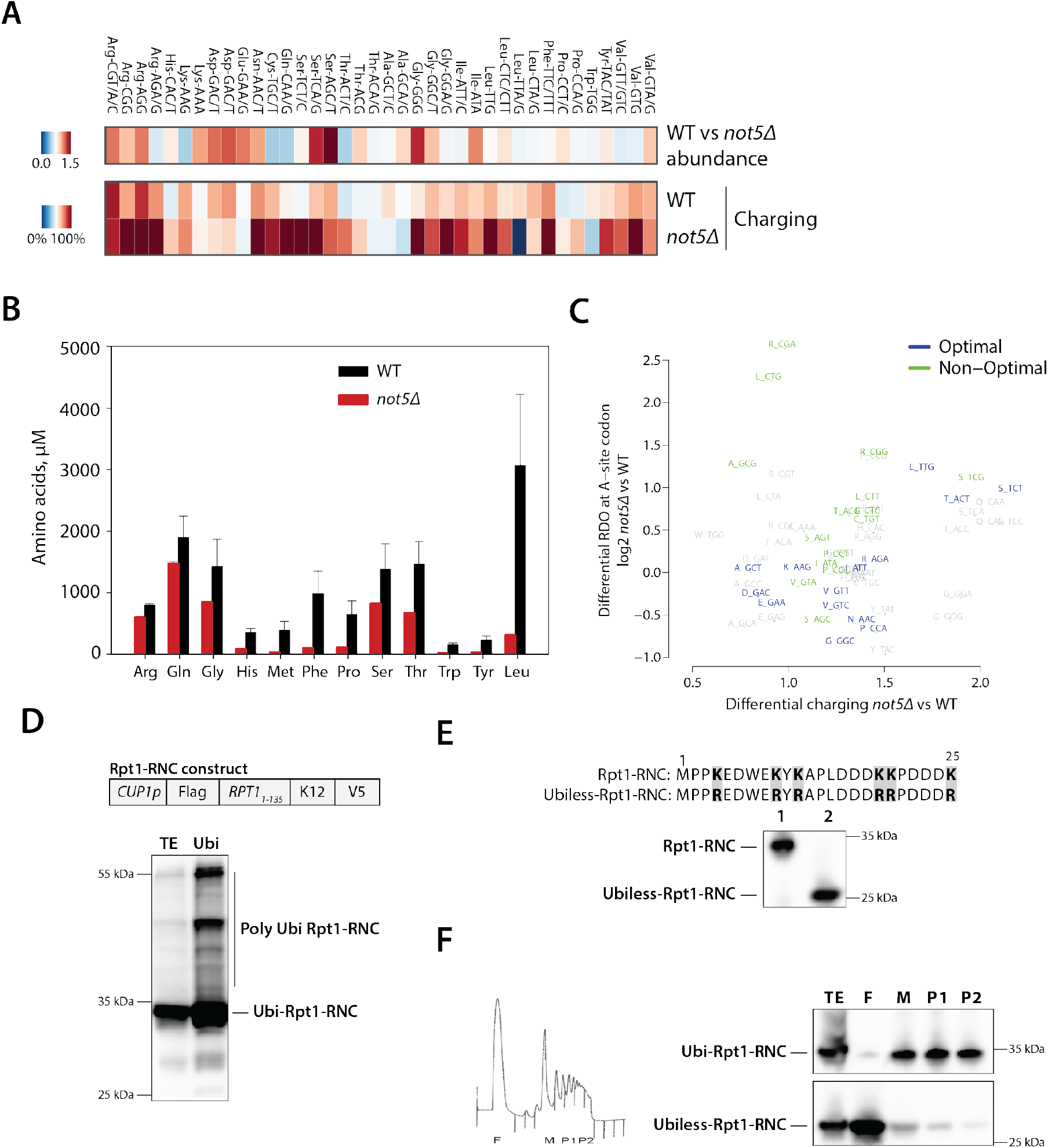
Not5 alters mainly elongation of highly abundant transcripts enriched with optimal codons. (**A**) Microarray of total (abundance) and aminoacyl–tRNAs (charging) in two independent biological replicates. tRNA probes are depicted with their cognate codon and the corresponding amino acid. (**B**) Metabolomic analysis of intracellular free amino acids. (**C**) Relative aminoacyl–tRNA charging levels (*not5Δ* vs wild type) for each cognate codon are plotted against the relative A-site RDO (log2FC *not5Δ* vs wild type) of these codons. Charging fold change was significantly increased in the higher RDO group (One-sided t-test, p = 0.006229; logFC>0.2) relative to lower (logFC<−0.2). (**D**) Ubiquitinated proteins were affinity purified from cells co-expressing a stalled nascent Rpt1 chain (RNC) (RNC construct above) and His6-tagged ubiquitin. The input extract (TE) or Ubi-affinity proteins (Ubi) were detected with antibodies to the N-terminal Flag tag. The discrete Rpt1-RNC is ubiquitinated (Ubi-Rpt1-RNC). Polyubiquitinated forms of the Rpt1-RNC are additionally visible (Poly Ubi Rpt1-RNC). (**E**) Expression of the Rpt1-RNC without (1) and with (2) the first 6 lysines mutated to arginine (mutations shown above) (Ubiless-Rpt1-RNC) monitored with anti-Flag antibodies. (**F**) Polysome profiling of cells expressing the Rpt1-RNC or the Ubiless Rpt1-RNC (as in E). Fractions were visualized with anti-Flag antibodies. F, free fractions (F), M, 80S monosomes, P1 light polysomes, P2, heavy polysomes (profile above).

Altered RDO at A-site codons in *not5Δ* increased the span between slow- and fast-translated codons. Consequently, it would increase the kinetic window for concomitantly occurring co-translational processes. Stalled ribosome nascent chain complexes (RNCs) are normally turned over by RQC and NGD but stabilized in Not1-containing granules (7). We determined that they are expressed in the form of ubiquitinated NCs of a discrete size (**Fig. 2D**). Preventing ubiquitination by mutating the first lysine residues of NCs (**Fig. 2E)** released the NCs from ribosomes (**Fig. 2F**) facilitating drop-off at stalled positions and aborting elongation, without reading through the stalling sequence to the stop codon (**Fig. S2A**). Stalled NCs in *not5Δ* were not all ubiquitinated (**Fig. S2B**), and the non-ubiquitinated NCs were released from the stalled ribosomes (**Fig. S2C**).

We observed earlier that Not5 deletion enhances protein aggregation (9), thus we reasoned that NCs released from abortive translation in *not5Δ* would aggregate. Proteomic analysis of the protein aggregates in *not5Δ* revealed that from the 192 proteins detected in the aggregates (threshold 2 or more peptides, **Table S3**), 143 were unique and absent in the aggregates of wild-type cells. Of those, 69.9% (p=2.2×10^−16^, Fisher’s exact test, **Table S4**) were amongst those aggregating in cells lacking the ribosome-associated Ssb chaperones (15), suggesting that proteins with chaperone-assisted folding have higher tendency to aggregate when translation kinetics is altered. We also noted that most of the proteins detected in the *not5Δ* aggregates were from highly expressed mRNAs (**Fig. S3A**), enriched in optimal codons and lower A-site RDO. Such mRNAs were also down-regulated in *not5Δ* (**Fig. 3A**).RP mRNAs, highly expressed and with a high bias towards optimal codons, showed a marked reduction of RPFs in *not5Δ* (**Fig. 3A,B**), with a major contribution to the reduced RPFs (**Fig. 3B**). However, reduced translation elongation in *not5Δ* was clearly detectable even after excluding the RP transcripts (**Fig. 3C**), supporting the general role of Not5 in elongation. Besides RP mRNAs, translation of several translation factors (e.g. Efb1, Yef3, Cam1, Dhh1, eIF5a) and chaperones (Ssz1, Zuo1, Ssb1) was down-regulated in *not5Δ* (**Fig. 3D** and **Table S1**). eIF5a, encoded by *HYP2* and *ANB1* in yeast, was amongst the most down regulated proteins, and its depletion caused a global translation elongation defect, phenotypically similar to that in *not5Δ* (**Fig. 3E**). However, eIF5a depletion did not reduce RP mRNA translation (**Fig. 3F**) or RDO at optimal codons; instead it displayed increased RDO at proline CCA codon, related to eIF5a function in facilitating elongation at proline codons (12,16). Comparison of RPF changes between wild type and *not5Δ* to those in newly synthesized proteins selected from a SILAC analysis (8) showed an overall good correlation (**Fig. 3D**), indicating that the detected RPF changes in *not5Δ* (**Fig. 1A**) originated mainly from changes in newly synthesized proteins.

**Fig. 3.**
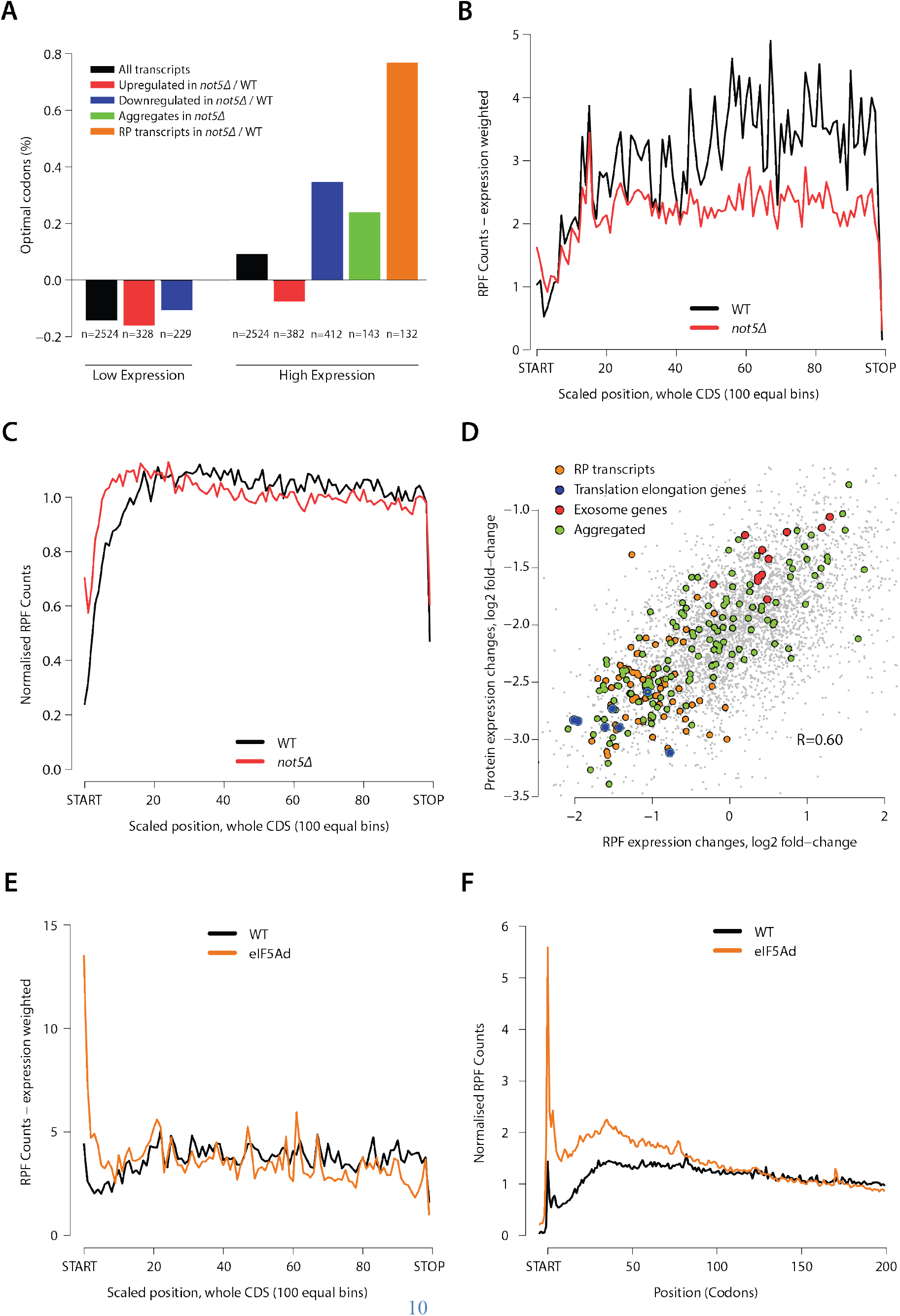
*not5Δ* translation elongation defect is distinct to that in eIF5A deficient cells. (**A**) Percentage of optimal codons (as in Fig. 1C) calculated for each CDS, averaged for various gene groups and log-normalized to the mean transcriptome-wide percentage. Low/high expression is defined as below/above median log RPKM. (**B**) Metagene RPF coverage of non-weighted RP mRNAs. Transcripts longer than 100bp are included. (**C**) Metagene analysis of all transcripts except RP mRNAs. (**D**) Comparison of normalized RPFs (ribosome profiling) and protein expression (SILAC) changes in *not5Δ* vs wild type. R, Pearson correlation coefficient. (**E, F**) Metagene analysis of eIF5A-degron strain for all transcripts (E) or of RPF coverage of non-weighted RP mRNAs (F).

The higher RDO at minor codons in *not5Δ*, indicative of ribosomal stalling, could trigger mRNA turnover. Supportive for this was the detected up-regulation in *not5Δ* of transcripts encoding 3’ to 5’ exosome with a role in co-translational quality control and no-go decay (NGD) (**Fig. 3D**). In contrast, the 5’ to 3’ exonuclease Xrn1, a key component of NGD (17), was down-regulated (8) and also found amongst the aggregated proteins in *not5Δ* (**Table S3**). The Dcp2-decapping factor was downregulated (8), whereas *Pat1*, which activates mRNA decapping in a Not5-dependent manner (18), was up-regulated. Likely, the upregulated *PAT1* and exosomal transcripts compensate for the decreased components of 5’ to 3’ degradation machinery in Not5 depleted background.

Together our findings indicate that Not4 and Not5 have distinct functions in translation, but collaborate at the transition from initiation to elongation (**Fig. 4**). In polysome profiles Not4 was mostly associated with monosomes, whereas Not5 was associated with both monosomes and polysomes (**Fig. S4A**). The deletion of Not5 reduced overall ribosomes in both monosomal and polysomal fraction (**Fig. S4B)**, while Not4 deletion reduced specifically only the polysomes (**Fig. S4C**). Deletion of Not5 increased the amount of Not4 in the monosomes (**Fig. S4D**) and Not4 deletion increased the levels of ribosomes co-purifying with Not5 (**Fig. S4E** and **Table S5**). Components of the 3’ to 5’ exosome and Dom34, which facilitates ribosome recycling (19), co-purified with Not5 but only in Not4-deleted background as revealed by mass spectrometry analysis of tandem-affinity purified Not5 (**Table S5**). It suggests that Not4 might play a role in turnover of Not5 bound to the ribosomes. Collaboration between Not4 and Not5 at translation initiation is also supported by the fact that the deletion of Not4 and Not5 is synthetically lethal (20).

**Fig. 4.**
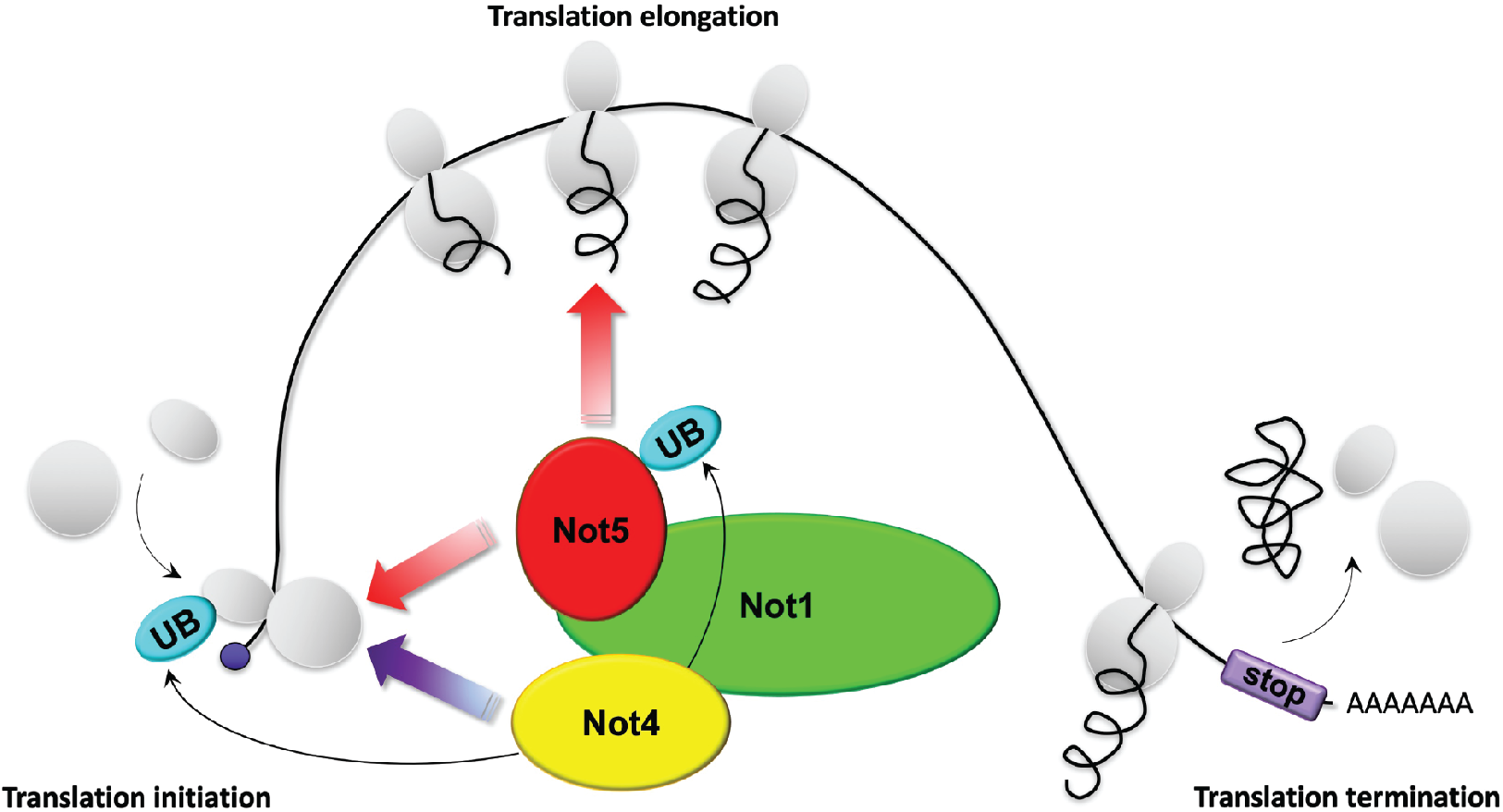
Not4 and Not5 regulate translation synchronously. Model of Not4 and Not5 collaboration in translation initiation and Not5 effect on elongation. Not4-dependent ubiquitination of Rps7 or other ribosomal proteins (9) or of Not5 (**Fig. S6C**) could facilitate the process.

Our data show that Not5 is important in translation elongation and that the overall reduced protein synthesis in *not5Δ* is a result of ribosomal drop-off mostly at mRNAs enriched in fast codons. *not4Δ* also showed codon-specific alterations in the RPF coverage of single transcripts, but the RDO changes were independent of codon optimality (**Fig. S5A**) and did not correlate with tRNA paucity (**Fig. S5B**) or free amino acid levels (**Fig. S5C**). The few observed changes in tRNA levels might reflect demand rather than be causal for the RDO alterations in *not4Δ*. The RDO changes unrelated to codon optimality in *not4Δ* had no effect on translation of RP mRNAs (**Fig. S5D**) but affected co-translational processes. For instance non-ubiquitinated NCs were also detected in *not4Δ* (**Fig. S6A**), released from ribosomes (**Fig. S6B**) and new proteins aggregate (21).

Based on our observations, we propose that Not5 and Not4 likely regulate the ribosome traffic for Not1-associated mRNAs to avoid downstream ribosomal collisions (**Fig. 4**), enabling Not1-containing assemblysomes to protect mRNAs with stalled ribosomes.

## Supporting information

Allen_Supplemental data

## Acknowledgments

We are grateful to Drs Ulrike Rolle-Kampczyk and Martin von Bergen (Helmholz for Environmental Research, Leipzig, Germany) for the metabolomics measurements.

## Author contributions

GEA performed ribosome profiling and SILAC analysis. OOP performed experimentally ribosome profiling and polysome fractionation experiments. ZV prepared the samples for aggregate, tRNA and metabolomics analyses. MZ did the *not4Δ* and *not5Δ* analysis of RNC expression and polysome fractionation. BW did RNA analyses and figures. CP performed and analyzed the tRNA microarrays, ZI and MC analyzed data, supervised the work and wrote the manuscript.

## Data and materials availability

tRNA microarray data are accessible under the accession number GSE137567 in the Gene Expression Omnibus (GEO) database and ribosome profililng of *not4Δ* and *not5Δ* GSE137613.

All code used in the manuscript is available https://github.com/georgeallenunige/NotTranslation

