## Supplementary material for "Switch from translation initiation to elongation needs Not4 and Not5 collaboration": Allen_Supplemental data

**This PDF file includes:**

Materials and Methods
Figs. S1 to S6
Table S4.
Captions for Data S1 to S4

**Other Supplementary Materials for this manuscript include the following:**

**Data S1**. Table S1. RPKM for the duplicate ribosome footprinting data samples of wild type *not4Δ* cells and *not5Δ* cells.

**Data S2**. Table S2. Microarray of total and aminoacyl-tRNA abundance in wild type *not4Δ* cells and *not5Δ* cells. The mean values were used in the heatmaps of Fig. 2A and Fig. S3C.

**Data S3**. Table S3. LC-MS analysis of aggregated proteins in *not5Δ* and wild type cells.

**Data S4**. Table S5. LC-MS analysis of proteins co-purifying with Not5-TAP from *not4Δ* and wild type cells.

### Materials and Methods

#### Strains, plasmids and oligonucleotides

Strains used in this manuscript were from either from Euroscarf and derived from BY4739 (*MAT*α *his3 leu2 ura3 lys2*): wild type (MY3415), *not4Δ* (isogenic except *not4::KANMX4*) and *not5Δ* (isogenic except *not5::KANMX4*) used for ribosome profiling, or the BY4742 background (*MAT*α *his3Δ leu2Δ lys2Δ0 ura3Δ*), wild type (MY3672), *not4Δ* (MY10776, isogenic except *not4::NATMX4*) and *not5Δ* (MY5673, isogenic except *not5::NATMX4*) or they have been described previously (Not4-TAP (MY4857) and Not5-TAP (MY5321)). TAP is a double affinity tag composed of the calmodulin binding peptide followed by a TEV (Tobacco Etch Virus) protease cleavage site and the Protein A tag (2). Not5-TAP *not4::NATMX4* (MY12006) and Not4-TAP *not5::NATMX4* ((MY5657)) were derived from MY5321 and MY4857 respectively (2). Plasmids for stalled nascent constructs were from pOP164 described previously (7) and obtained by PCR with oligonucelotides specific for different ORFs and cloning by gap repair. First six lysines of the Rpt1 ORF in pOP164 (pMAC1152) by PCR using an oligonucleotide bearing the mutations as a forward primer. The N-terminal truncated stalled Rpt1 (pMAC1149) has been described (7). All clones were verified by sequencing. Sequences of oligonucleotides are available upon request.

#### Ribosome profiling

Ribosome profiling of *not5Δ* (MY3418) and *not4Δ* (MY3417) was performed in biological duplicates as described (7). For the Ribo-Seq samples, all fastq files were adapter stripped using cutadapt (22). Only trimmed reads were retained, with a minimum length of 20 and a quality cutoff of 2 (parameters: -a 10 CTGTAGGCACCATCAATAGATCGGAAGAGCACACGTCTGAACTCCAGTCAC --trimmed-only --minimum-length=20 --quality-cutoff=2). Histograms were produced of ribosome footprint lengths that were very homogenous with highest reads between 28 and 31 that were kept for the analysis. Reads were mapped, using default parameters, with HISAT2 (23) to R64-1-1, using Ensembl release 84 gtf for transcript definitions. UTR definitions were taken from the Saccharomyces Genome Database and a standard region of 100bp was used where a gene's UTR was not defined. A minimum length of 30bp was implemented to ensure appropriate mapping around the start and stop codons. For the mapping, only unique alignments to transcripts were retained. A full set of 6692 CDSs were established for R64-1-1 Ensembl release 84 and extended by the same UTR sequences defined above. The filtered reads were then mapped to this transcriptome with bowtie2 (24), using default parameters.

For all downstream analysis, dubious ORFs were filtered to leave 5929 transcripts. The A/P-site position of each read was predicted by riboWaltz (25) and aggregated over all transcripts. Metagene plots were generated for genes with a mean of log2 RPKM above 5 (2604 for wild type and *not4Δ*, and 2849 for *not5Δ*). For the plots shown in Fig. 1A and Fig. 3E, genes were filtered if shorter than 600bp to ensure that metagene at start and stop did not overlap. For the scaled plot, genes longer than 100 nucleotides were retained and split into 100 equal bins. This excluded only 2 RP mRNAs. Normalization was performed as described (26).
Codon frequency was defined for each gene as the number of each codon type divided by the total number of codons in the CDS. The RDO was defined by the read count for each codon type divided by the total counts for the CDS and normalized to codon frequency. Mean RDOs for a group of genes was calculated as the average value for each codon. Genes used in the analysis of

RDO were restricted to those with RPKMs greater than 1 across all samples, leaving 5048 genes. The log2 ratio of *not4Δ* and *not5Δ* RDOs were taken relative to wild type for each codon. Differential expression was performed on the same set of genes (RPKM>1) comparing wild type and *not4Δ* or *not5Δ* duplicates using edgeR (27) on default settings. Where correlations are reported, the Pearson method was used.
To assess relative aminoacyl–tRNA charging levels for each cognate codon, where a codon is read by two tRNAs, charging was summed (weighted by its usage divided by the total codon usage of all codons read by each tRNA). Leu-TTA/G (fold change 0) was excluded through a Grubbs outlier test on charging fold changes ($p = 0.02956$).
Ribosome profiling data set of eIF5A-degron strain was reanalyzed from (12).

**Polysome fractionation.**
Cellular extracts were fractionated in sucrose gradients as described before (9). Proteins from the fractions were analyzed by western blotting.

**Protein aggregate analysis**

Protein aggregates were isolated as described previously (9) and a proteomic analysis performed by LC-MS at the core facility of the Faculty of Medicine (University of Geneva).

**Tandem affinity purification**

Not5-TAP was purified by tandem-affinity purification, first by IgG sepharose then by calmodulin affinity as described (2). Co-purifying proteins were identified by LC-MS.

***In vivo* ubiquitination assay**
The assay was performed as previously described (28) with cells expressing $His_6$-ubiquitin under the control of the *CUP1* promoter and ubiquitinated proteins were purified by nickel affinity chromatography.

**Antibodies**
Anti-Flag, V5, HA and PAP antibodies were commercial and anti-Rpl35 was a gift from Martin Pool.

**tRNA microarrays.** To determine the fraction of charged tRNAs we followed the procedure described in (14). Briefly, total RNA was isolated in mild acidic conditions (pH 4.5) which preserves the aminoacyl-moiety. Each sample was split into two aliquots and one was oxidized with periodate to which the changed tRNAs remain intact and following subsequent deacylation (100 mM Tris (pH 9.0) at 37°C for 45 min) was hybridized to Cy3-labeled RNA/DNA stem-loop oligonucleotide. The second aliquot was deacylated to receive the total tRNA and hybridized to Atto647-labeled RNA/DNA stem-loop oligonucleotide. Both aliquots were analyzed on the same tRNA microarrays with tRNA probes covering the full-length sequence of cytoplasmic tRNA species as described previously.

For tRNA abundance, total RNA was isolated at alkaline pH to simultaneously deacylate all tRNAs. tRNAs labeled with Cy3-labeled RNA/DNA stem-loop oligonucleotide were hybridized on the same microarray with tRNAs isolated from wild-type strain and labeled with Att647-labeled RNA/DNA stem-loop oligonucleotide. The arrays were normalized to spike-in standards, processed and quantified with in-house python and R scripts.

**Supplementary Figures**

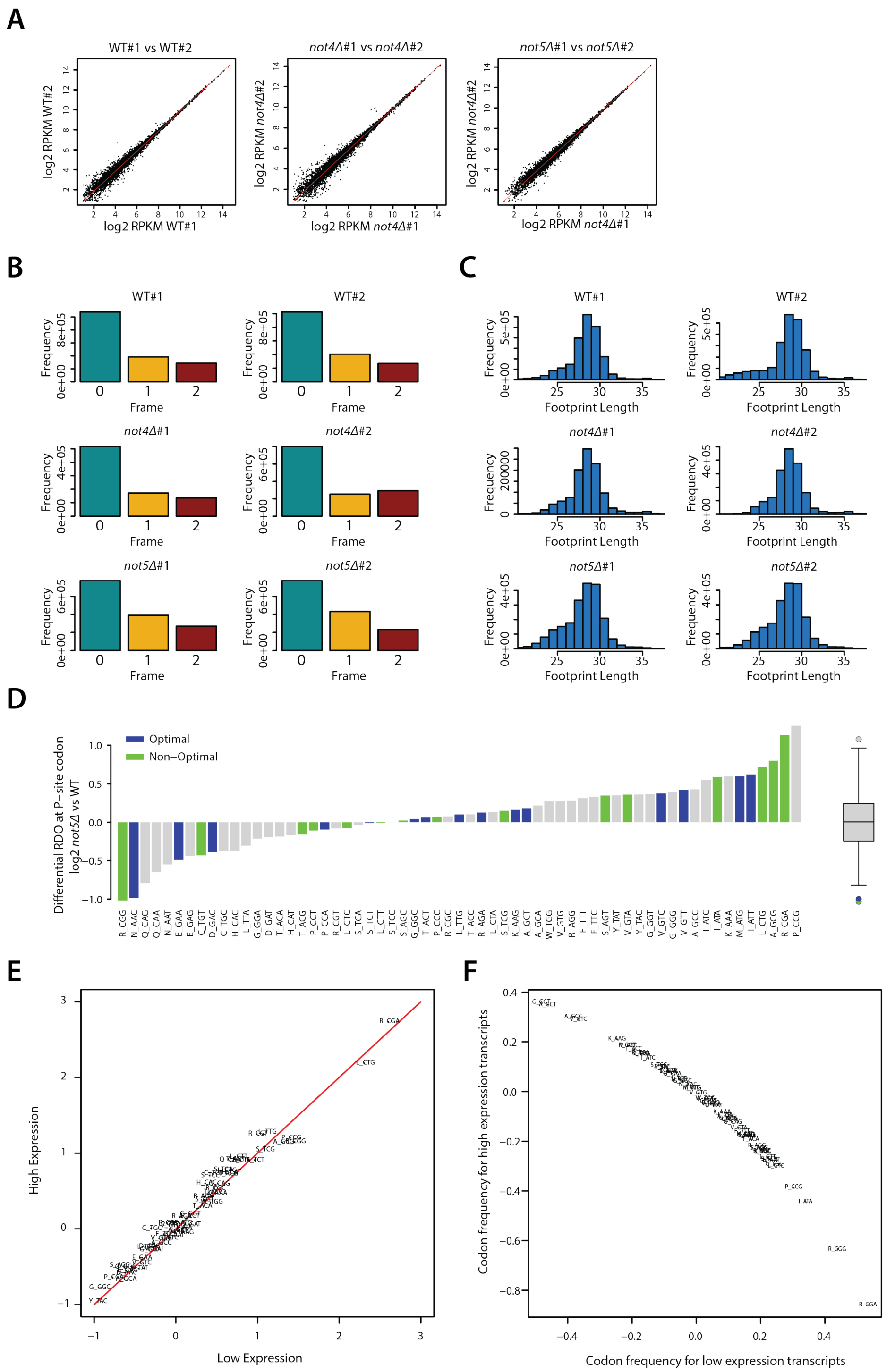


**Fig. S1**. **Quality control analysis of the ribosome profiling data.** (**A**). Comparison of the RPKM values for each transcript between biological replicates. (**B**) Three-nucleotide periodicity in the aggregated RPFs in each replicate. (**C**) Footprint length distribution. (**D**) RDO at P-site codons in *not5Δ* relative to wild type, with 15 most (blue) and 15 least used (green) codons. Grey, all other codons (**E**) Comparison of the RDO in wild type for the lower vs higher expressed halves of the 5048 transcripts (i.e. 2524 transcripts each half). (**F**) Codon frequency for the same two categories (as in E) normalized to the transcriptome codon usage.

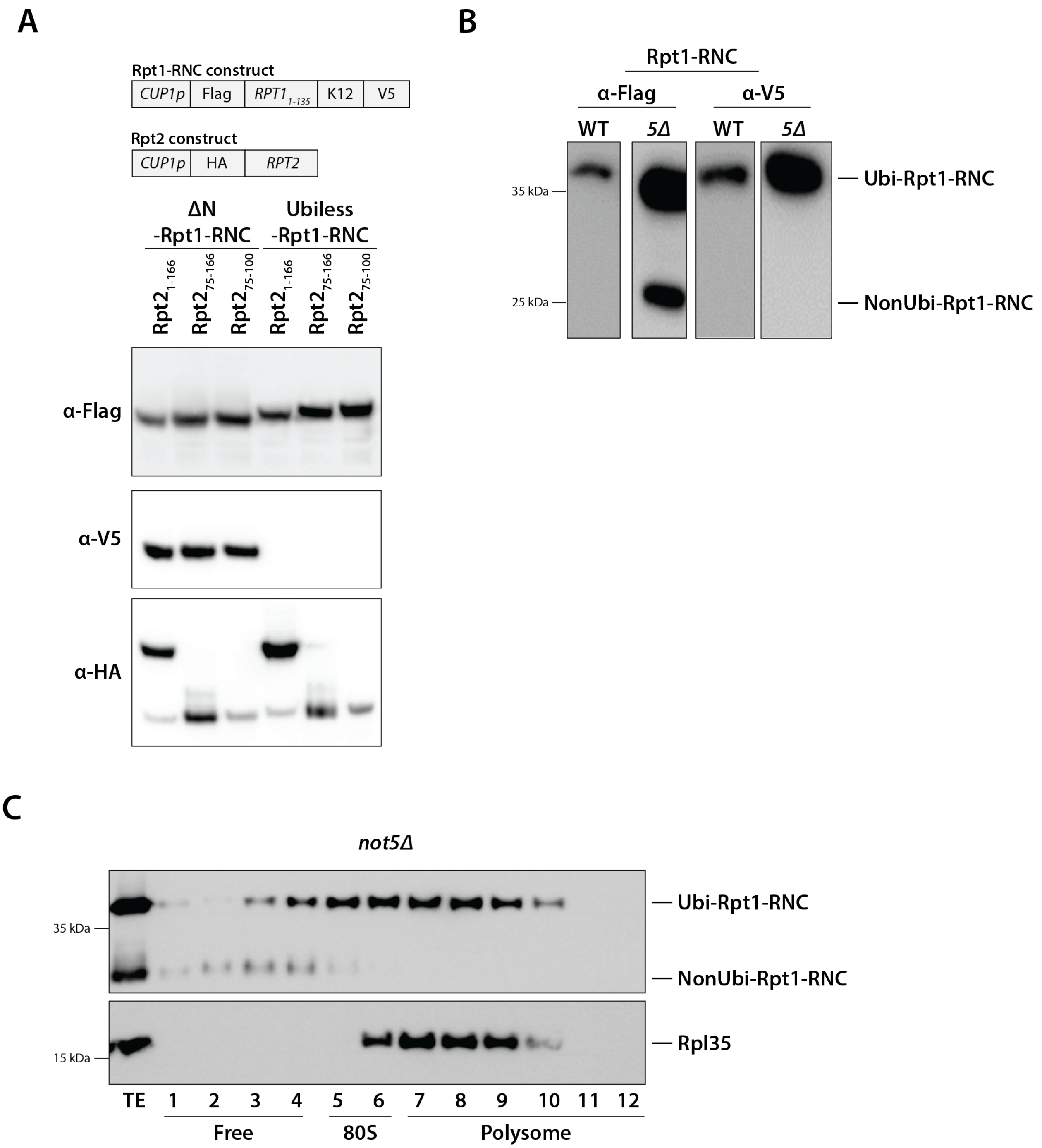


**Fig. S2. Loss of NC ubiquitination in *not5Δ* results in abortive translation following ribosome stalling.** (**A**) Cells co-expressing stalled Rpt1$_{49\text{-}135}$ nascent chain (7) (ΔN-Rpt1-RNC, first 3 lanes) or the Ubiless-Rpt1-RNC (last 3 lanes) together with different forms of the Rpt1 co-translational partner Rpt2 tagged at its N-terminus by HA were visualized with anti-Flag, V5 and HA antibodies (α–V5, α–Flag, α–HA), respectively. Constructs are depicted on the right. (**B**) Expression of the Rpt1-RNC in wild type or *not5Δ* (*5Δ*) was analyzed with anti-Flag and V5 antibodies as indicated. The ubiquitinated and non-ubiquitinated forms of the Rpt1-RNC are indicated. (**C**) Fractions from polysome profiles of same extracts analyzed in panel B were immunoblotted with anti-Flag and Rpl35, antibodies.

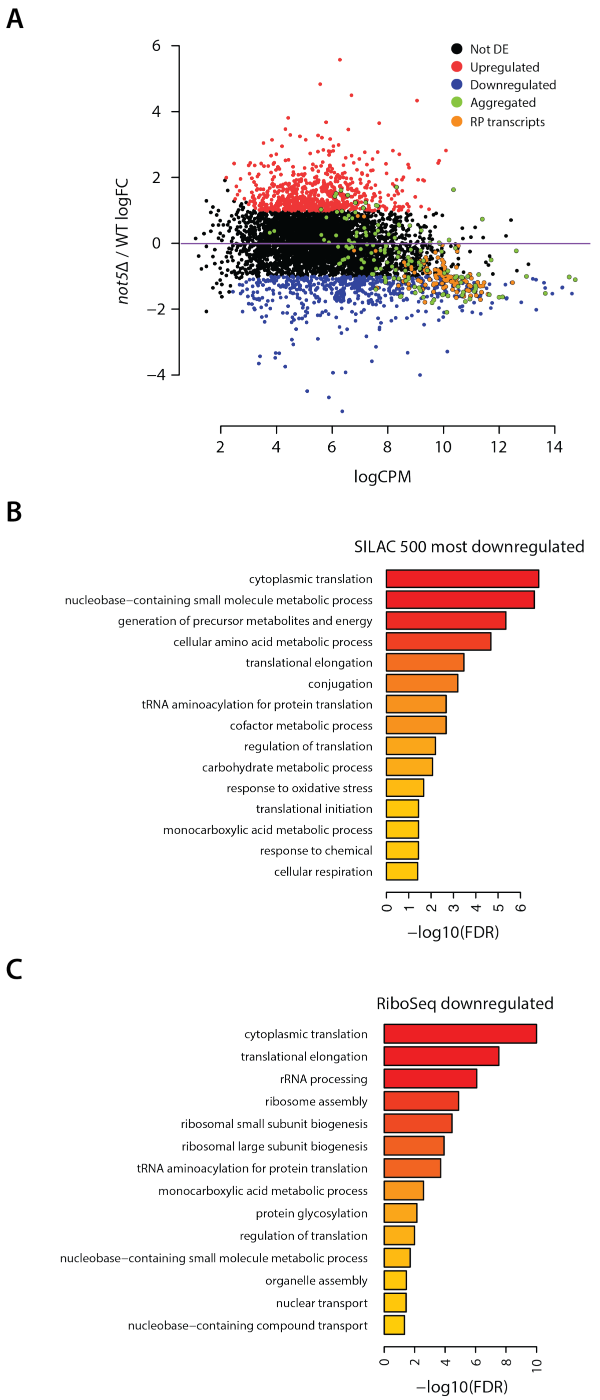


**Fig. S3.** (**A**) Comparison of the RPF profiles according to expression level of the wild type and *not5Δ*. Red, significantly up-regulated (FDR<0.05, logFC>1) transcripts; blue significantly down-regulated (FDR<0.05, logFC<-1) transcripts; green, transcripts producing aggregated proteins; orange, RP transcripts. (**B**) GO term analysis for the 500 most down-regulated proteins in *not5Δ* in SILAC (top) and ribosome profiling (bottom). FDR is calculated using Benjamini-Hochberg adjustment from p-values generated using a hypergeometric test.

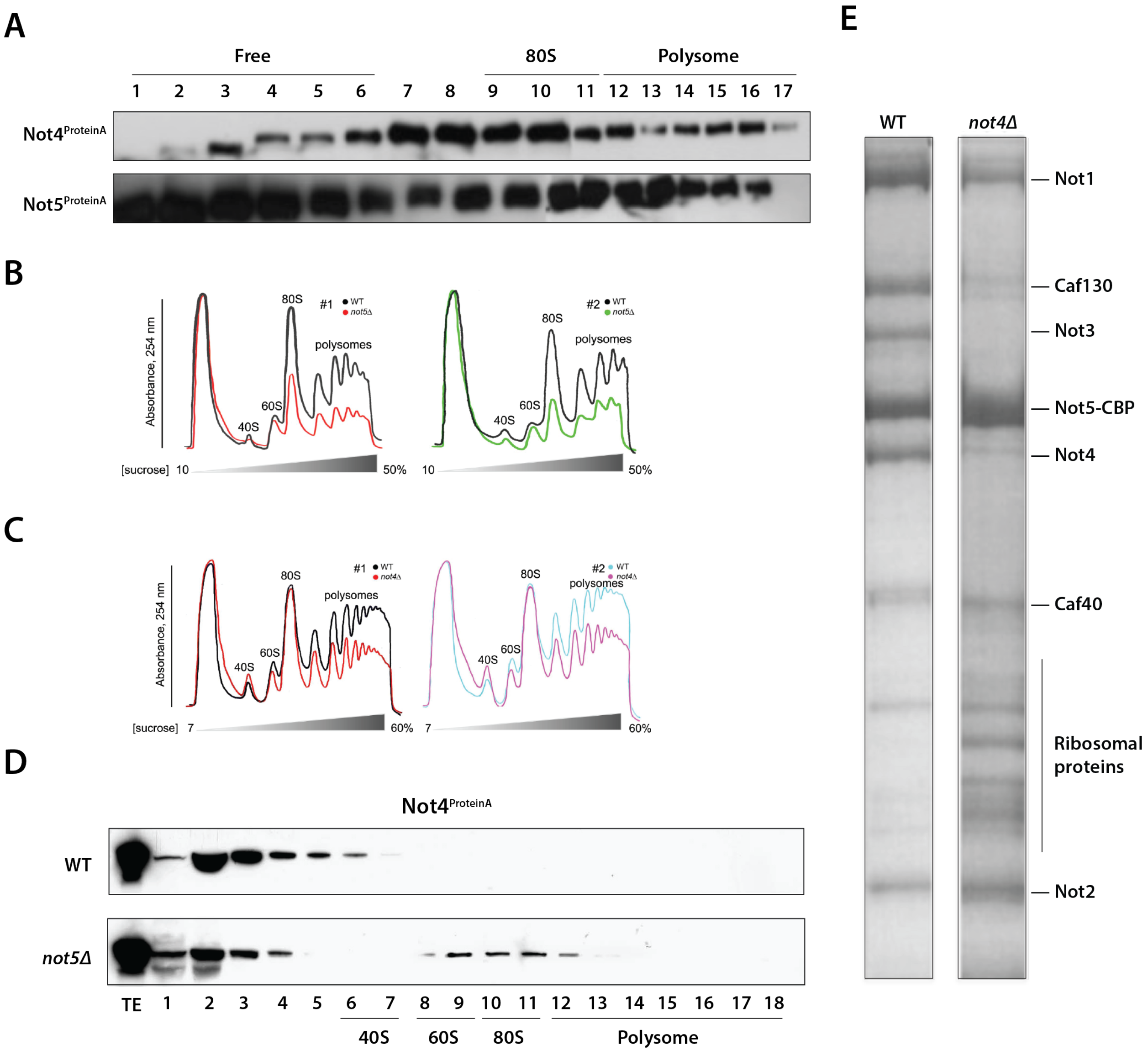


**Fig. S4. Not4 and Not5 associate with different ribosome pools**. (**A**) Wild-type cells expressing Not4 or Not5 fused to Protein A (Not4-ProtA, Not5-ProtA) were analyzed by polysome profiling and different fractions visualized with antibodies recognizing protein A (anti PAP). Free fractions (F), 80S monsomes and polysomes are indicated. (**B, C)**. Polysome profiles obtained by sucrose gradient sedimentation (7-50% or 7-60% as indicated) from two biological replicates of *not5Δ* (B) and *not4Δ* (C) overlaid with those from wild type. (**D**) Wild-type cells and *not5Δ* cells expressing Not4-ProtA were analyzed by polysome profiling followed by immunostaining of the different fractions with anti-PAP antibodies. (**E**) Proteins co-purified with Not5-TAP in 2 sequential affinity steps, from wild type or *not4Δ* cells, were separated by SDS-PAGE and visualized by Coomassie staining. Each gel lane was cut in slices and analyzed by LC-MS (**Table S5**). The position of the subunits of the Ccr4-Not complex and some ribosomal proteins is indicated. After tandem affinity Not5 remains fused to the calmodulin binding protein moiety (CBP) and is indicated (Not5-CBP).

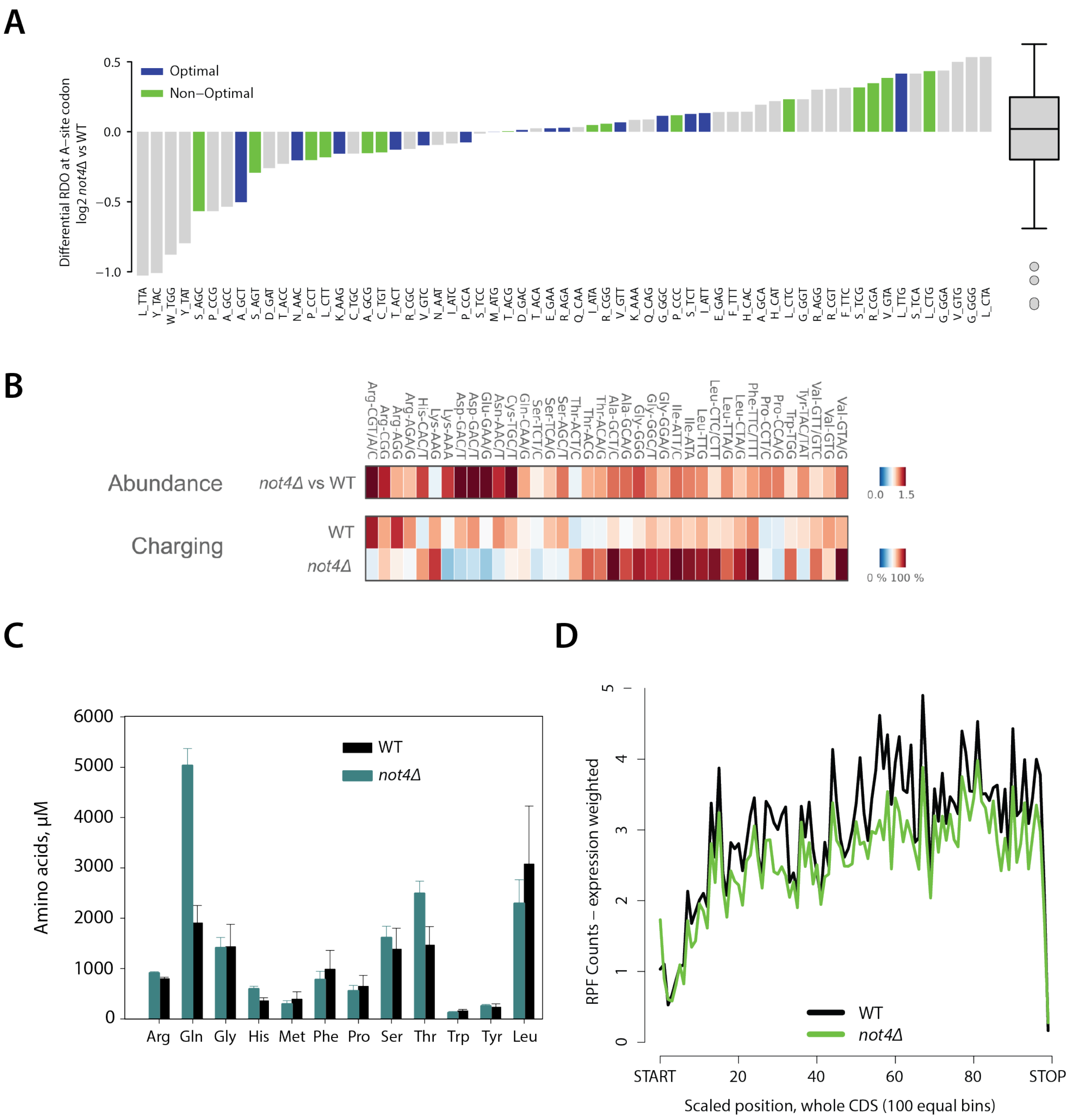


**Fig. S5. Ribosome profiling in *not4Δ* does not correlate with changes in tRNAs, amino acids or codon optimality.** (**A**) RDO at A-site codons in *not4Δ* relative to wild type, with 15 most (blue) and 15 least used (green) codons. Grey, all other codons. (**B**) Microarray of total (abundance, top panel) and aminoacyl–tRNAs (charging, bottom panel) represented as a mean value from two independent biological replicates. tRNA probes are depicted with their cognate codon and the corresponding amino acid. (**C**) Metabolomic analysis of free amino acids in *not4Δ*. (**D**) Metagene analysis of RP mRNAs in *not4Δ* (same analysis as for *not5Δ*, Fig. 3B).

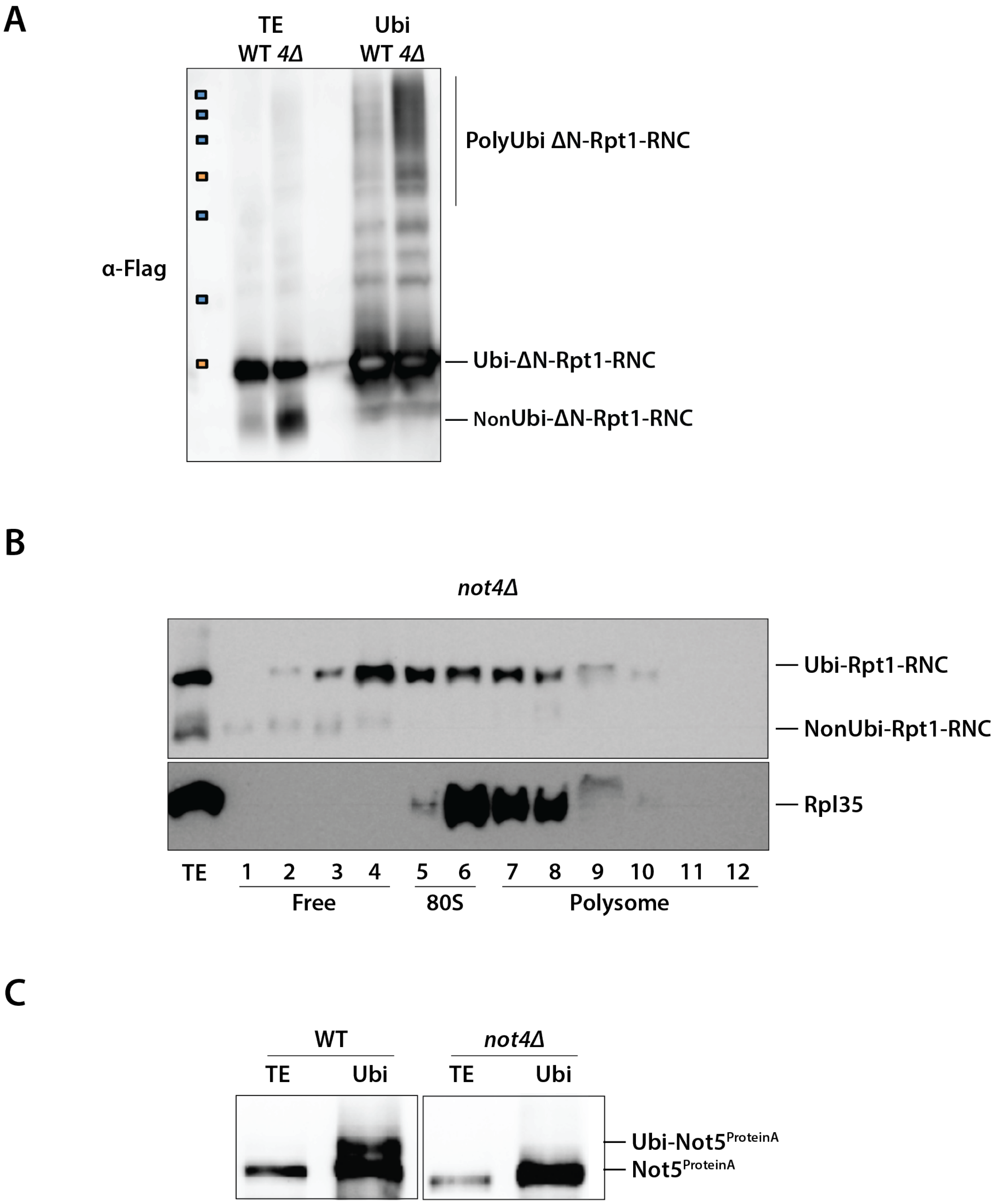


**Fig. S6. Not4 affects NC and Not5 ubiquitination.** (**A**) Ubiquitinated proteins from wild type or *not4Δ* cells expressing ΔN-Rpt1-RNC were affinity purified and analyzed as in Fig. 2D. (**B**) Fractions from polysome profiles of *not4Δ* cells expressing Rpt1-RNC analyzed by immunostaining with anti-Flag or anti-Rpl35 antibodies, respectively. (**C**) Ubiquitinated proteins from wild type or *not4Δ* cells expressing Not5-ProtA (from its own locus) and plasmid-borne ubiquitin were affinity purified and analyzed as in panel A. Total input extract (TE) or Ubi proteins visualized by immunostaining with anti-PAP antibodies.

| | |
|---|---|
| Total of proteins found in wt aggregates | 45 |
| Total of proteins found in not5Δ aggregates | 175 |
| Total of proteins found in ssbΔ aggregates[15] | 752 |
| Total of proteins found in not5Δ aggregates but absent in wt | 143 |
| Total of proteins found in not5Δ and ssbΔ aggregates but absent in wt | 100 |
| Total of proteins found both in wt and not5Δ | 32 |
| Total of proteins found in wt, not5Δ and ssbΔ | 24 |
| % Proteins found in not5Δ but not in wt that are found in ssbΔ | 69.9 |

**Table S4. Comparison or protein aggregates in *not5Δ* and cells lacking Ssb proteins.**
